# Neural network models for the evolution of associative learning

**DOI:** 10.1101/2023.07.21.549996

**Authors:** Emiliano Méndez Salinas, Franz J. Weissing, Magdalena Kozielska

## Abstract

The ability to learn from past experience is an important adaptation, but how natural selection shapes learning is not well understood. Here, we investigate the evolution of associative learning (the association of stimuli with rewards) by a modelling approach that is based on the evolution of neural networks (NNs) underlying learning. Individuals employ their genetically encoded NN to solve a learning task with fitness consequences. NNs inducing more efficient learning have a selective advantage and spread in the population. We show that in a simple learning task, the evolved NNs, even those with very simple architecture, outperform well-studied associative learning rules, such as the Rescorla-Wagner rule. During their evolutionary trajectory, NNs often pass a transitional stage where they functionally resemble Rescorla-Wagner learning, but further evolution shapes them to approximate the theoretically optimal learning rule. Networks with a simple architecture evolve much faster and tend to outperform their more complex counterparts on a shorter-term perspective. Also, on a longer-term perspective network complexity is not a reliable indicator of evolved network performance. These conclusions change somewhat when the learning task is more challenging. Then the performance of many evolved networks is not better than that of the Rescorla-Wagner rule; only some of the more complex networks reach a performance level close to the optimal Bayesian learning rule. In conclusion, we show that the mechanisms underlying learning influence the outcome of evolution. A neural-network approach allows for more flexibility and a wider set of evolutionary outcomes than most analytical studies, while at the same time, it provides a relatively straightforward and intuitive framework to study the learning process.

## 1. Introduction

The ability to learn from past experience is an important adaptation and learning capabilities have been identified in most animals and even in unicellular and non-animal species [1–5]. However, still very little is known about the evolution of learning.

From an evolutionary perspective, learning is an intrinsically difficult phenomenon to study because it encompasses two intermingled levels of adaptation: adaptive changes of cognition and behaviour during individual lifetime, and selection-induced changes of the mechanisms underlying learning over the generations. Furthermore, learning is a complex process involving many mechanisms (perception, processing of information, decision-making, emotions or other evaluation systems), which are only partly known and likely species-specific to a considerable extent. Last, but not least, it is not easy to judge the fitness consequences of a given learning mechanism, as the implications of this mechanism for survival and reproduction depend strongly on both the particular life trajectories of individuals (which determine what can and will be learned) and on the spatiotemporal structure of the environment.

It is therefore not surprising that only relatively few attempts have been made to model the evolution of learning. The most prominent approach considers the evolution of ‘learning rules’ [6–11]. The idea is that the learning process of an individual is characterized by a function *f* that describes how the cognitive state or behaviour *E*_*t*_ of an individual is updated on the basis of new information *R* on the individual’s environment or performance: *E*_*t*+1_= *f* (*R, E*_*t*_). The learning rule *f* is typically assumed to be characterized by a small number of heritable parameters (e.g. the slope and intercept determining a linear function) that evolve through natural selection, as individuals with more efficient learning rules transmit the underlying parameters to a larger number of offspring. We will later illustrate this approach by considering the evolution of the parameters characterizing the “Rescorla-Wagner rule” [12], which plays a prominent role in theories of associative learning. While the learning rule approach is relatively simple and elegant, it has the important drawback that the evolvable learning processes are strongly constrained from the start (by the limitations imposed on the set of learning rules), thus often preventing the evolution of more efficient ways of learning from experience.

To remove this restriction, Trimmer and colleagues [13] studied the evolution of associative learning by a more flexible genetic programming approach. They implemented the update process underlying learning by binary trees of arbitrary complexity. Such trees can not only implement simple learning rules but also highly intricate learning processes. Moreover, a binary tree provides an easy way to implement inheritance of learning mechanisms (including mutations and recombination). While in case of parameterized learning rules the evolutionary outcome can often be predicted by analytical methods (by optimality or game theoretical techniques), this is no longer possible for genetic learning programmes; instead individual-based simulations are required to predict the course and outcome of evolution. Based on such simulations, Trimmer *et al*. [13] concluded that learning programmes resembling the Rescorla-Wagner rule emerge in the early stages of evolution, but that these are later replaced by more efficient programmes. The genetic programming approach has the big advantage that it allows the evolution of a broad range of learning processes, including highly efficient processes that would often be excluded *a priori* from the set of learning rules considered by the modeller. On the downside, the evolved genetic learning programmes are often difficult to interpret, as programmes for similar or even identical learning processes can have a very different structure (see Appendix D in [13]). Moreover, the encoding of the genetic programmes by binary trees and, more importantly, their inheritance and mutation, is quite artificial and non-intuitive from a biological and behavioural perspective.

Here, we use a different approach by considering the evolution of neural networks (NNs) that underly associative learning. NNs allow for the implementation of a similarly broad range of learning strategies as the genetic programming approach. As neural networks play a crucial role in organismal learning, modelling learning using NNs have the added advantage that knowledge of the neuronal basis of learning can be included in the model in a natural manner. Here, however, we do not attempt to replicate the structure and function of any real biological neural system. Instead, we employ NNs with a very simple architecture as a conceptual tool for understanding how learning can evolve in a system that, just as the neural networks of real organisms do, receives and processes information from the environment to update their internal states which in turn can modify future responses to the environment.

We consider the evolution of NNs that have to solve a simple associative learning task, which has fitness consequences for the learning individual. The parameters determining the NN are genetically encoded and transmitted from parents to their offspring. NNs inducing more efficient learning have a selective advantage and spread in the population. Over the generations, selection will shape NNs which, for a given network architecture, perform the learning task increasingly well. By systematically comparing different network architectures, we will address the following research questions: To what extent is NN-based learning comparable to existing learning paradigms, such as the Rescorla-Wagner rule? How well do evolved NNs perform in comparison to the theoretically optimal performance in the given learning task? What is the relationship between network complexity and network performance?

On purpose, we focus on NNs with a very simple architecture, as only simple NNs allow the systematic analysis of the simulation results with the standard toolbox of evolutionary theory (e.g. the analysis of fitness landscapes). By increasing network complexity in a step-by-step manner, we hope to build up knowledge and expertise that will be useful when analysing more complex networks. For our study, we will make use of the simple learning task designed by Trimmer and colleagues [13]. This way, we can use their results as a benchmark for judging the learning performance of evolved NNs.

## 2. Associative learning from an evolutionary perspective

Associative learning, the phenomenon that organisms come to associate different stimuli or events with each other, is widespread among animals and arguably the most important paradigm in the study of learning [14]. This form of learning has obvious evolutionary implications as it allows organisms to use formerly neutral stimuli (known as conditioned stimuli) to predict the presence/arrival of fitness relevant events (known as unconditioned stimuli). For these reasons, studying a simple associative learning task is a good starting point for investigating the evolution of learning.

Take an imaginary example of a bumblebee that visits flowers of different colours in a meadow for the first time. It does not know which type of flower provides the best source of nectar. By visiting flowers of a specific colour, say purple, the bumblebee might learn that this colour is indicative of a rich nectar reward. By correctly associating purple colour with nectar presence, a bumblebee could spot “nectary” flowers by their purple colour from a far distance and, in this way, forage more efficiently in the meadow, thus enjoying a clear fitness advantage.

The current modelling convention for associative learning in the field of experimental psychology is to predict the changes in the associative strength (usually measured from the frequency/strength of a behavioural responses observed in experiments) between two given stimuli. We will here follow Trimmer *et al*. [13], who reinterpreted “associative strength” as an estimate of the probability of receiving a reward. Continuing with our example, every time the bumblebee visits a purple flower, it will update its estimate of the probability of finding nectar in this type of flowers. If in our hypothetical meadow the probability that purple flowers offer nectar is high, a bumblebee will be rewarded with nectar most of the times it visits a purple flower. Accordingly, it should associate purple flowers with a high probability of finding nectar. Should our bumblebee move to another meadow where purple-flowering plants offer no nectar, it should reduce its estimate of the probability that purple flowers provide nectar with each non-rewarded visit.

The Rescorla-Wagner model [12] is arguably the most influential model of associative learning [15,16]. Trimmer and colleagues [13] posed the question whether, why, and when the “Rescorla-Wagner rule” would evolve through natural selection. To this end, they took the simplest form of this model [17], which describes the build-up of an association between a single conditioned and unconditioned stimulus. Interpreting the association strength between the two stimuli as the subjective probability of receiving a reward, this version of the Rescorla-Wagner rule can be written as:

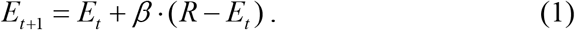

This rule describes how the estimate *E*_*t*_ of the reward probability is updated after receiving an input *R*, which can take on the values *R* = 1 or *R* = 0 depending on whether or not a reward was present. The parameter *β* determines the learning rate. Trimmer *et al*. [13] compared the performance of evolved genetic learning programmes with this version of the Rescorla-Wagner rule. We will follow their lead and do the same for evolved learning networks.

## 3. The model

### 3.1 Model overview

We consider a population of individuals that are faced with the task of determining the true state *P* of their local environment on the basis of learning experiences, where *P* corresponds to the probability of receiving a reward. Each individual is endowed with a heritable neural network, which processes incoming information to sequentially update an estimate *E* of the true state *P*. We assume that the reproductive success of an individual is inversely related to the estimation error *δ* ^2^ = (*E* − *P*)^2^. Hence, individuals with a small discrepancy between their learned estimate *E* and the true value *P* produce more offspring, which inherit their parent’s network, subject to rare mutations. Over the generations, networks will evolve an improved ability to estimate the true state of their environment based on the information provided in this environment.

### 3.2 The learning task and the effect of learning on reproductive success

Following Trimmer *et al*. (2012), we consider the associative learning task illustrated in Figure 1. During its lifetime, each individual is confronted with *K* different environments (grey rectangles) that differ in their probability *P*_*k*_ of obtaining a reward. When entering a new environment, the individual starts with the same initial estimate *E*_0_ of the reward probability. Subsequently, the individual has *S* learning experiences. In experience *t*, the individual either receives a reward or not (*R*_*t*_ = 1 or 0), where *R*_*t*_ is randomly drawn from a binomial distribution with success probability *P*_*k*_. Based on its learning network, the individual uses the value *R*_*t*_ to update its estimate of the reward probability from *E*_*t*_ to *E*_*t* +1_. After *S* learning experiences, a final estimate *E*_S_ is obtained. The squared difference 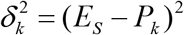 between the estimated reward probability *E*_*S*_ and the true reward probability *P*_*k*_ quantifies the estimation error made in environment *k*. The total lifetime error ∆_*i*_ of individual *i* is the average of its *K* errors. The reproductive success of individual *i* is proportional to 1− (∆_*i*_ ∆_max_), where ∆_max_ is the maximal lifetime error made in the population. Throughout, following Trimmer *et al*. [13], we chose *K* = 6. The reward probabilities *P*_*k*_ were drawn independently from a uniform distribution U(0,1). At first, we consider a simple learning task in which the number of learning updates is fixed (*S* = 10) and later a more complicated tasks with variable *S*.

**Figure 1:**
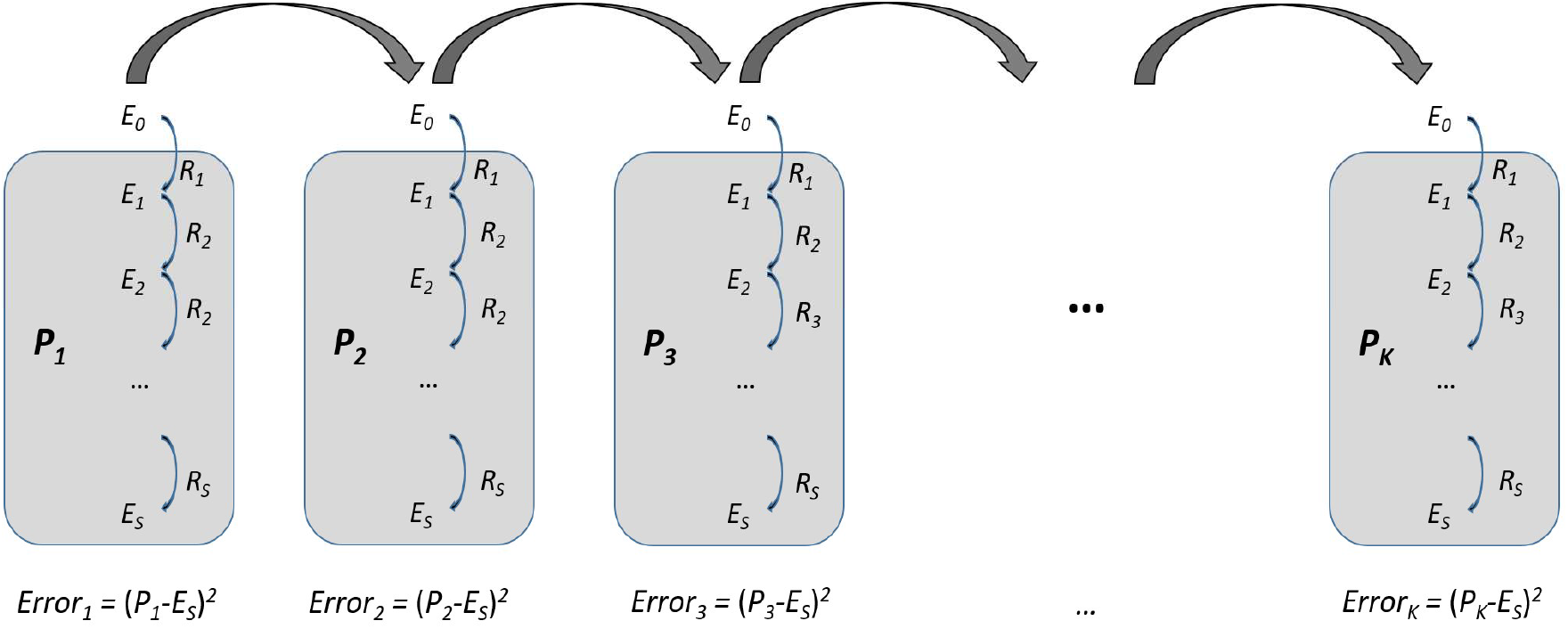
Overview of the learning task. Individuals are faced with *K* different environments, each with a different probability of reward *P*_*k*_. In each environment the individual has *S* learning experiences after which its estimation error is calculated (see text for details).

### 3.3 Network structure

In our study we consider simple neural networks (NNs) [18], like those in Figure 2. In the standard version, our network receives two inputs, *E*_*t*_ (the current estimate of the reward probability) and *R* (the presence or absence of a reward), and it produces an updated estimate of the reward probability, *E*_*t* +1_, as its single output. The networks consist of nodes (the circles in Figure 2) that are organized in a sequence of layers. Each node is connected to one or several nodes in the subsequent layer (the arrows in Figure 2), and it can stimulate or inhibit the activities of these nodes. Each connection has a certain weight *w*, where a positive value of *w* represents stimulation, while a negative value corresponds to inhibition. The nodes of the input layer take on real values that correspond to either *E*_*t*_ or the reward *R* received (where *R* = 0 or *R* = 1). These values are processed and determine the node activities at the subsequent level. More precisely, the activity the form *y*_*i*_ of node *i* in the consecutive layer is given by an expression of the form

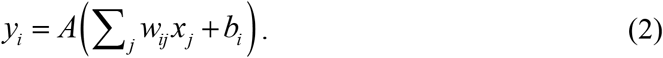

**Figure 2:**
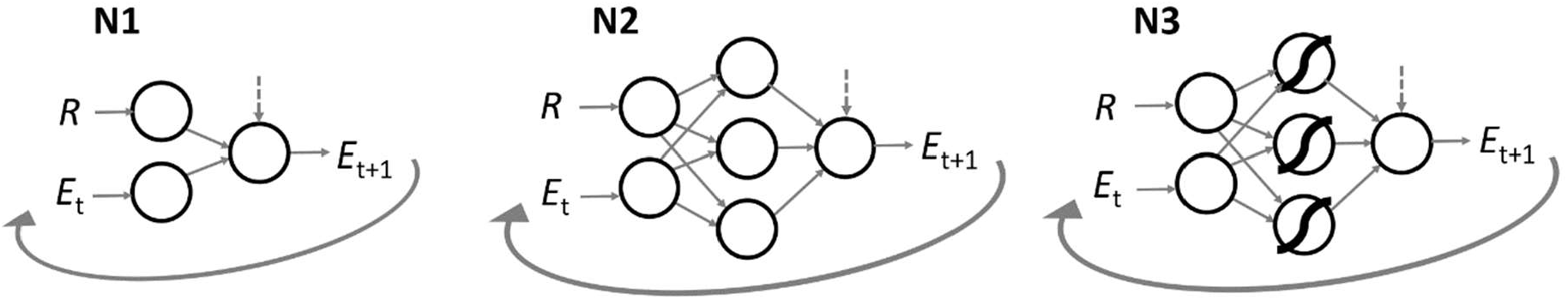
The three basic network architectures studied. **N1:** Single-layer network with linear transmission of information. **N2**: Two-layer network with linear transmission of information. **N3**: Two-layer network with non-linear activation function in the processing layer. Vertical dashed arrows indicate that a bias is added to the output. We also analysed the same kind of networks in the absence of an output bias; we will refer to these networks as **N1**_**0**_, **N2**_**0**_ and **N3**_**0**_, respectively. In the first learning step, *E*_*t*_ is given by a genetically determined initial estimate *E*_0_. In each subsequent learning step, the output *E*_*t* +1_ of the previous step becomes the input *E*_*t*_ of the new step.

Here *j* runs over all nodes of the previous layer that are connected to *i, x*_*j*_ is the activity of node *j*, and *w*_*ij*_ is the strength of the connection between nodes *j* and *i. b*_*i*_ is a so-called “bias” that corresponds to the baseline activation of node *i*. Finally, the function *A* is a so-called “activation function.” Such functions can be useful, as they allow for more versatile input-output relationships of an NN and because they can ensure that the activity levels *y*_*i*_ are restricted to a certain range (such as the interval [-1,+1]).

An NN and, hence, the iterative process *E*_*t* +1_ = *f* (*R, E*_*t*_) induced by it, is fully determined by its architecture (i.e. the nodes and their connection patterns), the activation functions, and the values of the connection strengths *w*_*ij*_ and the biases *b*_*i*_. As described in Section 3.4, we assume that the network parameters *w*_*ij*_ and *b*_*i*_ are heritable and transmitted from parents to offspring.

In our study, we will investigate the evolution of the networks **N1, N2** and **N3** in Figure 2 in considerable detail. Networks of type **N1** are particularly simple in that they do not have a processing layer and because the activation function is just the identity function. This implies that the estimate of the reward probability is updated according to the linear relationship

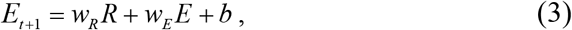

where the heritable (and hence evolvable) parameters *w*_*R*_, *w*_*E*_, and *b* correspond to the weights of the connections from *R* to *E*_*t* +1_ and from *E*_*t*_ to *E*_*t* +1_ and to the bias of the output node *E*_*t* +1_, respectively. Networks of type **N2** have also the identity function as their activation function, but they include a 3-node processing layer. This implies that they also encode a linear input-output relationship while having more evolutionary degrees of freedom (see Appendix A). In networks of type **N3**, the activation function of the nodes in the processing layer is non-linear. We considered various options for the activation function, but we here only report the results on the hyperbolic tangent function (*A*(*z*) = tanh(*z*)), a widely used S-shaped function with output values in the interval (−1,1), which also performed best in our preliminary studies.

In the second part of our study (where the number of learning updates *S* is variable), we will also consider networks which have an additional input that is related to the number of updates experienced so far. This way, the NNs could potentially use this kind of information to modulate their learning over time.

### 3.4 Reproduction and inheritance

In our simulations, we consider a population with discrete, non-overlapping generations, where each generation consists of *N* =1,000 haploid individuals. Each individual harbours genes that encode the connection weights and biases of its NN and the initial estimate *E*_0_ of the reward probability. These genes can take any real value. In a network of type **N1**, for example, there are four genes that encode the values *w*_*R*_, *w*_*E*_, *b* and *E*_0_. The alleles of an individual at these gene loci determine how the network of the individual functions, i.e., how the reward probability *E* is updated by the presence or absence of a reward (see eq. (3)).

When all individuals of a given generation have experienced their *K* environments, their reproductive success is determined as described in Section 3.2. The population reproduces and is replaced by the population of their offspring. For each of the *N* offspring, a parental individual is drawn at random, with probability that is proportional to the individual’s reproductive success. The offspring inherits all network parameters from its parent. With per-locus mutation probability *µ* = 0.0015, a parental allele is affected by a mutation. In such a case, a small number ε is added to the parental value, where the mutational step size ε is drawn from a normal distribution with mean 0 and standard deviation 0.01.

Most simulations were run for 500K (= 500,000) generations. In case of the networks considered by us in detail (Figure 2), an evolutionary equilibrium was usually reached in a much shorter time.

### 3.5 Network performance and optimally performing NNs

Our individual-based simulations are subject to stochasticity from different sources, like random mutations, limited population size, the random choice of *K* environmental parameters *P*, and the reward sequence in each environment (i.e., the sequence of reward/no-reward outcomes encountered by an individual during the learning process). To overcome the effects of this stochasticity, we quantify the “performance” of an evolved NN by the *expected* error **E**(*δ* ^2^) made by the network when estimating the reward probability of an environment. This corresponds to the error to be expected given a large number of environments and reward sequences as explained below.

As above *δ* ^2^ = (*Es* − *P*)^2^ indicates the squared deviation between the final estimate *E*_*S*_ and the true reward probability *P*, and **E**(*δ* ^2^) is the squared deviation to be expected in a random learning trial (after *S* learning events). Notice that **E**(*δ* ^2^) is an “inverse” performance measure, as a high network performance is associated with a low value of **E**(*δ* ^2^). For the linear networks **N1** and **N2** analytical expressions for **E**(*δ* ^2^), both for fixed and variable number of updates *S*, are derived in Appendix A. This allows to determine the parameters of the *optimally* performing NN by minimizing **E**(*δ* ^2^).

With the same procedure, we can also determine the performance measure **E**(*δ* ^2^) for learning rules such as the Rescorla-Wagner rule or the rule for Bayesian updating. Analytical expressions for **E**(*δ* ^2^) could not be found for networks of type **N3** with a non-linear activation function, therefore they were derived numerically (see Appendix A for details). Throughout, we will use **E**(*δ* ^2^) when comparing NNs with each other or with specific learning rules.

### 3.6 Visualisation of results by Rescorla-Wagner plots

According to equation (1), the Rescorla-Wagner rule is characterized by ∆*E* = *E*_*t* +1_ − *E*_*t*_ = *β* (*R* − *E*_*t*_). In other words, plotting the change Δ*E* in association strength against *R* − *E*_*t*_ results in a straight line through the origin, with a slope that is given by the learning rate *β*. In order to compare the learning behaviour of our evolved NNs with Rescorla-Wagner learning, we use the NN to calculate ∆*E* = *E*_*t* +1_ − *E*_*t*_ for a range of *E*_*t*_ values and both options *R* = 0 and *R* = 1 ; subsequently, ∆*E* is plotted against *R* − *E*_*t*_. We will call these plots “Rescorla-Wagner plots”. Such plots provide an intuitive way to visualise how different NNs behave in comparison to each other and in comparison to the Rescorla-Wagner rule and other learning rules (see also Appendix B).

## 4. Results

### 4.1 Simple learning task

In order to get a firm understanding of network evolution, we will first analyse a baseline scenario with a fixed number of *S* = 10 updates and an initial estimate *E*_*0*_ = 0.5 that is fixed and not evolvable. In Section 4.2, we will later investigate a more complex task, where the number of updates is not specified beforehand.

#### 4.1.1 Learning rules

For this baseline scenario, the optimal value of the learning parameter in the Rescorla-Wagner rule is *β* ^*^ = 0.135 (see Appendix A and [13]). In other words, given the reward *R*, the optimal updating of information according to the Rescorla-Wagner rule is of the form:

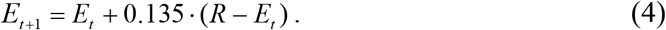

As shown by Trimmer *et al*. [13], there is, however, a learning rule (which they call the ‘optimally performing rule’ or OPR) that is as simple as the Rescorla-Wagner rule but, in the baseline scenario, provides on average a better estimate *E*_*S*_ of *P* than the Rescorla-Wagner rule:

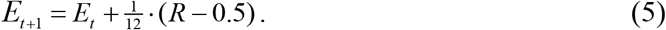

After *S* = 10 updates, this rule is equivalent to Bayesian updating, and thus represents the best performance possible under this scenario.

#### 4.1.2 Evolved network performance in comparison to learning rules

All our evolving reference networks, **N1** to **N3** (with and without bias), are able to outperform the RW rule to different degrees. The two linear networks without bias (**N1**_**0**_ and **N2**_**0**_) are able to achieve only a slightly better performance level than the best Rescorla-Wagner rule, while the two linear networks with bias on the output node (**N1** and **N2**) approximate the performance of the optimally performing rule OPR (Figure 3). The non-linear network without bias (**N3**_**0**_) is able to reach a better performance than the RW rule (Appendix C, Figure A2), but this happens only in a few replicates, and it takes a very long time. The non-linear network with bias (**N3**) approaches the performance of ten rounds of Bayesian updating, but this never happens within the timeframe of 500K generations used in our simulations (Figure 3). In general, non-linear networks perform worse than their linear equivalents.

**Figure 3:**
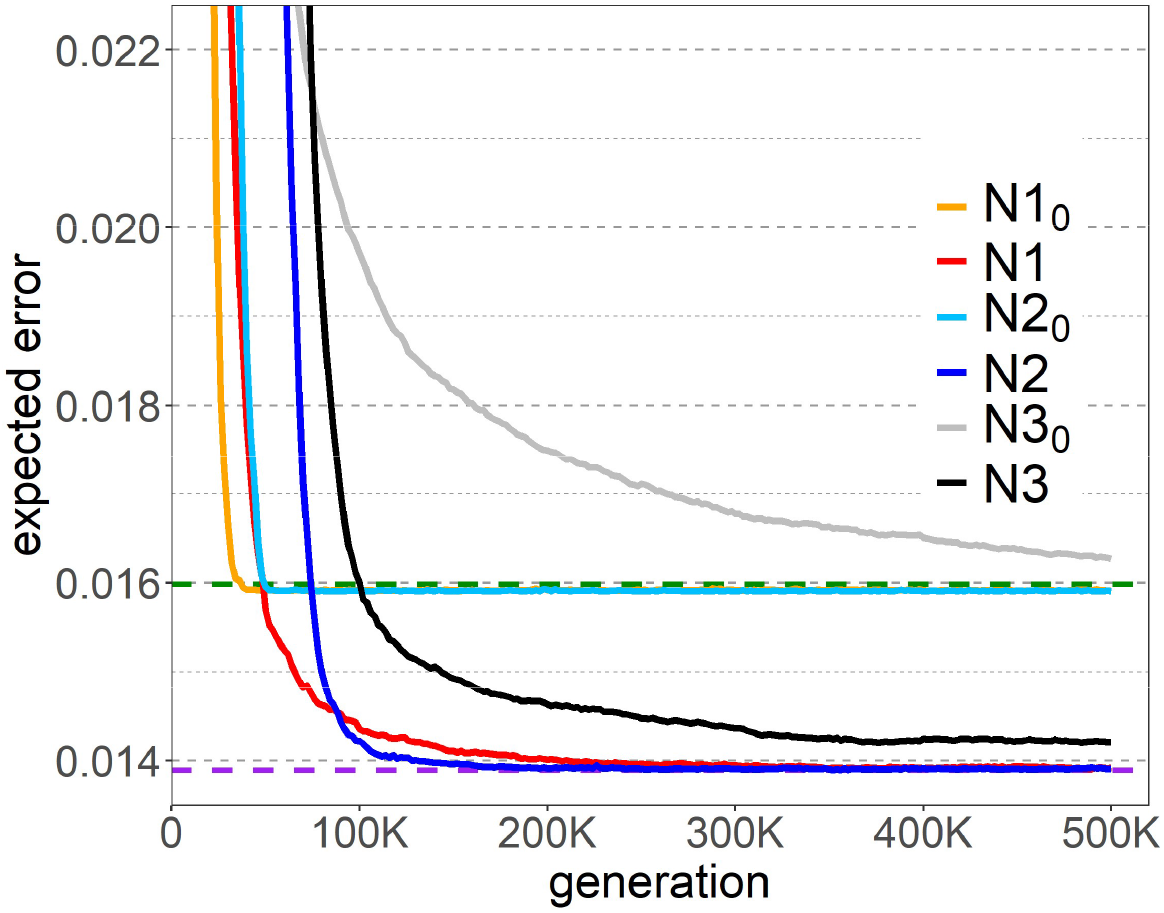
Performance of evolved networks in the simple learning task and *E*_0_=0.5. The graph shows for each type of network how the error decreases (and hence performance increases) over the generations. For comparison, the expected error of the Rescorla-Wagner rule (green dashed line) and the Optimally Performing Rule (purple dashed line) are also shown. Only networks with bias (**N1, N2**, and **N3**) clearly outperform the RW rule, with **N1** and **N2** reaching optimal performance. Linear networks without bias (**N1**_**0**_ and **N2**_**0**_), marginally outperform the RW rule. Networks of type **N3**_**0**_ can outperform the RW rule, but this only happens in small fraction of the replicates, and it takes a long evolutionary time (see Appendix B, Figure A2). Each solid line in the graph shows the average expected error from 10 replicate simulations, where each replicate is represented by best-performing NN in the population at the given time.

We will now have a closer look at the evolutionary trajectories of the four linear networks. A network of type **N1**_**0**_ is characterized by two evolvable parameters *w*_*R*_ and *w*_*E*_, and the estimate of the reward probability is updated according to the linear relationship that is a special case of equation (3) with *b*=0:

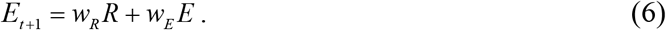

Notice that Rescorla-Wagner updating (1) corresponds to the special case *w*_*R*_ + *w*_*E*_ = 1 (where *w*_*R*_ = *β* and *w*_*E*_ = 1− *β*). Since a network of type **N1**_**0**_ allows for more general updating (two free parameters instead of one) one might expect that networks can evolve that outperform the best Rescorla-Wagner rule (3). However, mathematical analysis (see Appendix A) shows that the optimal network of type **N1**_**0**_ uses the updating algorithm

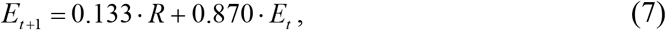

which is very close to the optimal RW rule (4) and reduces the expected error of that rule by only 0.5% (see Table A1, Appendix A). The **N1**_**0**_ networks in our individual-based simulations evolved weights and performance levels that are very similar to the mathematical optimum.

As networks of type **N1**_**0**_ have only two evolvable parameters, their evolution can be visualized on the corresponding fitness landscape. Figure A3 (Appendix D) shows that the single fitness peak is very close to the intersection point of the ‘line of attraction’ and the line *w*_*R*_ + *w*_*E*_ = 1 characterizing all those **N1**_**0**_ networks that are equivalent to a RW rule. This explains our observation in Fig. 3 that networks of type **N1**_**0**_ only achieve a marginally better performance than the RW rule, although they have two, rather than one, evolvable parameters. Even though optimal networks of type N1_0_ can slightly outperform the Rescorla-Wagner rule in our baseline scenario, they were indistinguishable from it when looking at the average population performance in the individual-based simulation.

A network of type **N1** has a bias *b* on its output node, leading to the update rule given by eq. (3). Only networks with such a bias can match the performance of the optimally performing rule OPR, because a bias is needed to produce the free term 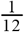·(−0.5) in eq. (5). Mathematical analysis reveals that the optimal updating of a **N1** network is indeed given by eq. (5), which characterizes the OPR (see Table A1, Appendix A). Networks of type **N1** do indeed converge to the OPR, both in their weighing factors and in performance (Figure 3).

For networks with non-linear activation function, **N3** and **N3**_**0**_, we were not able to find the optimal weights (or weight combinations) analytically.

#### 4.1.3 Learning behaviour of evolved networks

Trimmer et al. [13] based many of their conclusions on the comparison of performance of different binary trees, and not on the actual learning behaviour (they only analysed few binary trees in detail). For example, they implicitly assumed that binary trees that match the performance of a RW rule show similar updating behaviour as the RW rule. However, this is not necessarily the case. To get an impression of the updating behaviour of the evolved NNs, we produced Rescorla-Wagner plots for the evolved networks (see Section 3.6), as these plots reveal congruences as well as differences in updating behaviour in comparison to both the RW rule and the optimally performing rule (see also Appendix B).

Figure 4 shows that – for the simple learning scenario with *S*=10 and *E*_0_ = 0.5 – networks that perform like the RW rule or the OPR also tend to show the same (or a very similar) updating behaviour as the corresponding rule (but see section 4.2.2 for a different outcome): Evolved networks of type **N1**_**0**_ and **N2**_**0**_ exhibit practically the same updating behaviour as the best RW rule; and evolved networks of type **N1** and **N2** match the updating behaviour of the optimally performing rule. The behaviour of the non-linear networks (**N3**_**0**_ and **N3**) resembles the learning behaviour of the rules they approximate in performance, but only for a limited range of input values.

**Figure 4:**
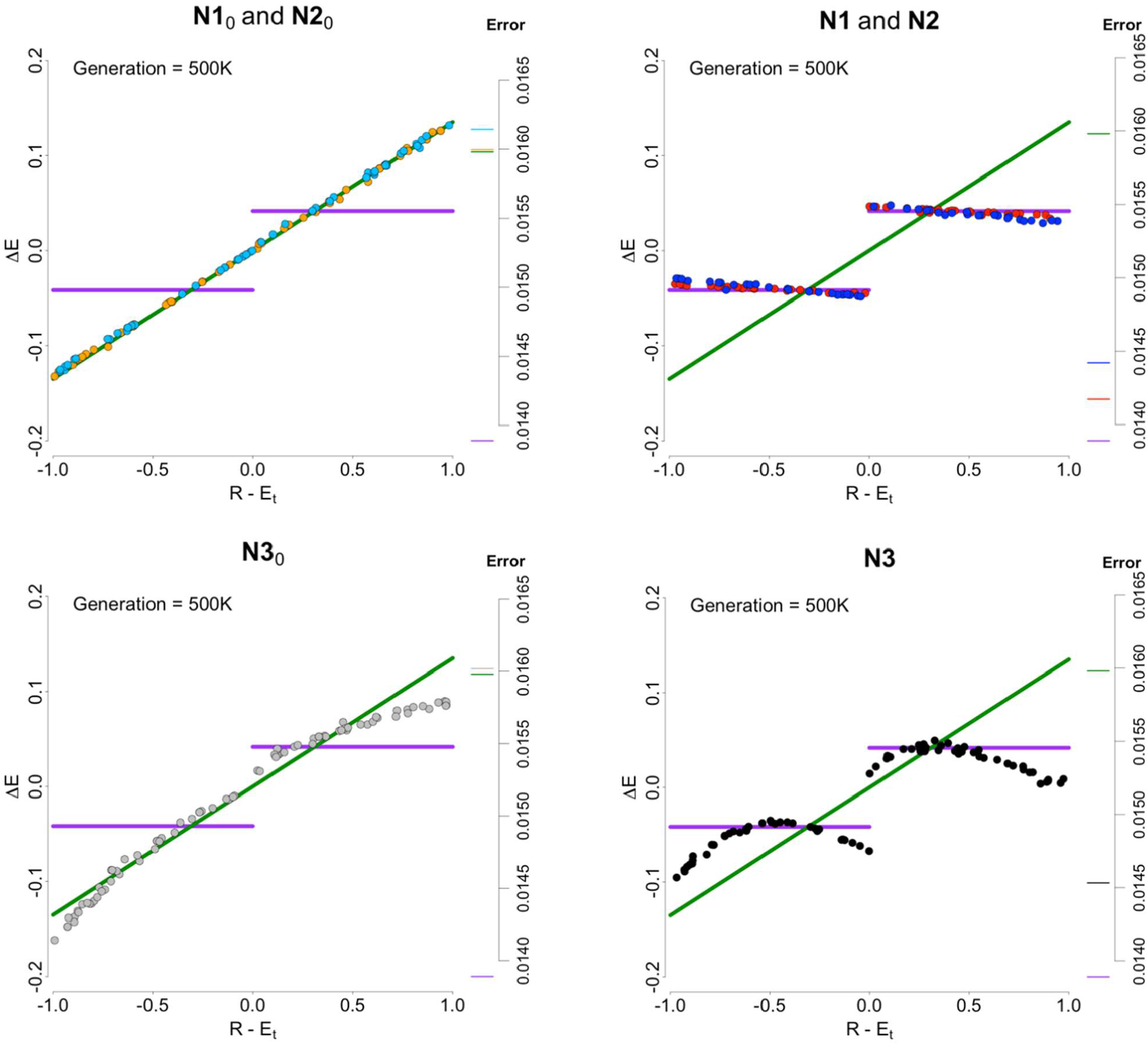
Rescorla-Wagner plots of NNs that evolved in the simple learning scenario. When the change in estimates (Δ*E* = *E*_*t* +1_ − *E*_*t*_) in response to an observation *R* (*R* = 0 or *R* = 1) is plotted against the difference *R* − *E*_*t*_, for a range of values 0 ≤ *E*_*t*_ ≤ 1, the Rescorla-Wagner rule (eq. (1)) produces a straight line with slope *β* through the origin. The green line corresponds to the best Rescorla-Wagner rule (*β*\* = 0.135). The optimally performing rule OPR (eq. (4)) produces two horizontal line segments (purple). The panels show the updating behaviour of evolved NNs of the six network types considered. In line with their performance (Figure 3), the updating behaviour of linear networks is very close to that of the RW rule when the networks do not have bias (**N1**_**0**_ : orange dots, **N2**_**0**_ : light blue dots) and very close to that of the OPR when they have a bias (**N1** : red dots, **N2** : blue dots). The updating behaviour of the non-linear networks corresponds to a concave function that only partly matches the RW rule (**N3**_**0**_ : grey dots) or the OPR (**N3** : black dots).

As shown in Figure 5, **N1** and **N2** networks ultimately match the updating behaviour of the OPR. Interestingly, these networks show Rescorla-Wagner kind of updating behaviour at an intermediate stage of their evolution (see Appendix E, Figure A4). Similarly, Trimmer *et al*. [13] reported that many of their genetic programmes performed similarly to the Rescorla-Wagner rule in an initial stage of evolution, but that these programmes were eventually replaced by programmes outperforming this rule. Our analysis of network behaviour shows that they not only resemble RW rule in performance but also in behaviour.

**Figure 5:**
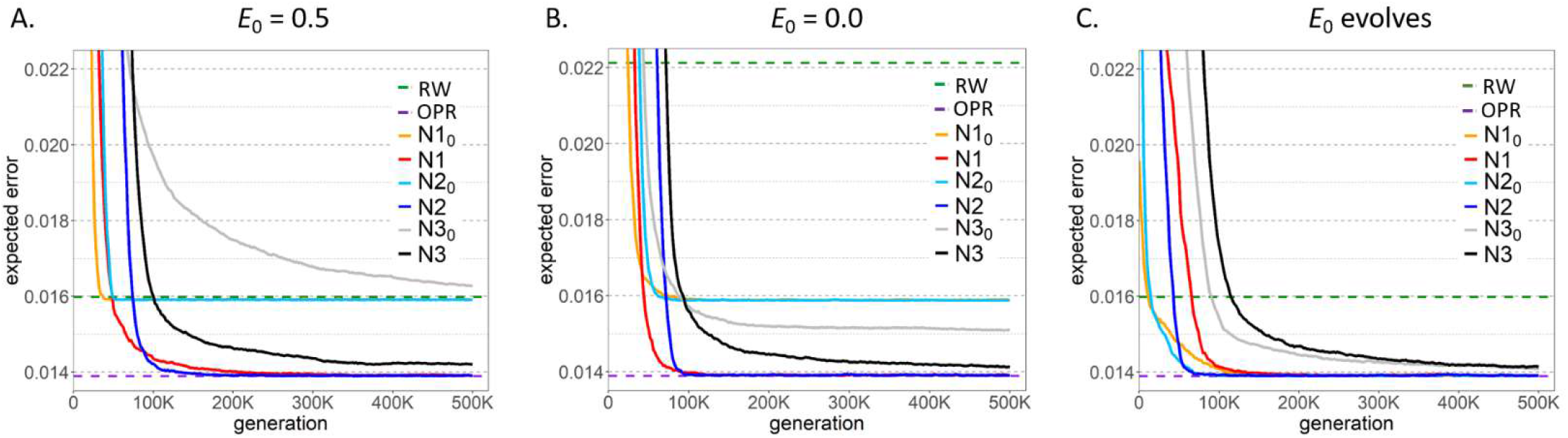
Effect of *E*_0_ on the performance of evolved NNs (simple learning task). The solid lines show the evolutionary increase in performance (= the decrease in expected error) of six types of network for three scenarios for the initial estimate *E*0 of the reward probability: **(A)** *E*0 = 0.5, **(B)** *E*0 = 0.0, **(C)** *E*0 jointly evolves with the other network parameters. The dashed lines indicate the performance of the best Rescorla-Wagner rule (green) and the Optimally Performing Rule (purple). For ease of comparison, panel (A) is a replicate of Fig. 3. Panel (B) illustrates lower performance (= higher error) of the RW rule when *E*0 = 0.0, as discussed in Section 4.1.4. In this case, all networks outperform the best RW rule, but only the linear networks with bias (**N1** and **N2**) achieve optimal performance. Panel (C) shows that all networks approach optimal performance if *E*0 is allowed to evolve. All graphical conventions are as in Figure 3.

#### 4.1.4 E_0_ and the interpretation of the Rescorla-Wagner rule

Until now we considered a situation in which the initial estimate of the probability of receiving a reward is fixed to *E*0 = 0.5. When *E*_*t*_ is interpreted as an estimate of the probability of receiving a reward, a prior of 0.5 makes sense, as in our model the reward probabilities are drawn from a uniform distribution over the unit interval. However, when the traditional interpretation of Rescorla-Wagner rule is used, *E*_0_ = 0.5 does not seem to be the most obvious initial condition. In the classical conditioning experiments that led to the development of the RW rule, the conditioned stimulus (CS) is, at the start of the experiment, usually new and neutral for the subject. In other words, there is initially no association between different stimuli or stimuli and response. Therefore, an initial value *E*_0_ = 0 seems the most obvious choice when *E*_*t*_ is interpreted as the association strength between stimulus and reward.

Interestingly, in our model the performance of the Rescorla-Wagner rule is much poorer for *E*_0_ = 0 than for *E*_0_ = 0.5, even if the learning rate *β* is optimized separately for both cases (the optimal learning rate is *β*\* = 0.196 if *E*_0_ = 0 and *β*\* = 0.135 if *E*_0_ = 0.5). This conclusion can be drawn from mathematical analysis, which shows that, quite generally, the performance of the best RW rule decreases in our model with the distance of *E*_0_ to 0.5. This is confirmed by individual-based simulations: when both *β* and *E*_0_ evolve, *E*_0_ converges to 0.5, and *β* converges to 0.135.The most likely explanation lies in the fact that, in our model, the RW rule is shaped by natural selection and adapted to a situation where the conditioned stimulus is not arbitrary (as in a typical learning experiment) but drawn from a fitness-relevant distribution. In other words, the organism can already have some “inherited” knowledge of the structure of the environment. It would be interesting to see whether the time course of learning in an associative learning experiment will differ depending on whether the conditioned stimulus is arbitrary or relevant for fitness.

#### 4.1.5 Effect of *E*_0_ on network evolution

Figure 5 shows that, in contrast to what we observed for the Rescorla-Wagner rule, the neural networks evolved the same level of performance, irrespective of whether the initial estimate of the reward probability was *E*_0_ = 0.5 or *E*_0_ = 0 (Fig. 5AB). The initial value *E*_0_ = 0 even seems to be slightly superior: it speeds up evolution and networks of type **N3**_**0**_ achieve a better performance. We conclude that network evolution is less sensitive to the initial estimate than optimizing the RW rule, most likely because networks have more evolutionary degrees of freedom.

As shown in Figure 5C, network evolution is speeded up and network performance is generally improved when *E*_0_ is a heritable trait that is jointly evolving with the network weights and biases. This holds especially for the networks without bias (**N1**_**0**_, **N2**_**0**_, and **N3**_**0**_) that now approach optimal performance, while this was not the case when *E*_0_ was fixed. Upon closer inspection, these networks evolved *E*_0_ values close to 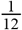, allowing them to perform almost as well as the OPR and the networks with bias. The networks with a bias already achieved (**N1, N2**) or approached (**N3**) optimal performance for a fixed value of *E*_0_ (Fig. 5AB). One might have expected that for these networks *E*_0_ would evolve to the value 0.5, the natural prior in our model. As shown in Section 4.4.1, this does indeed happen in case of the RW rule, where *E*_0_ converges to 0.5 when *E*_0_ can jointly evolve with the learning rate *β*. However, this did not happen in the networks with a bias (**N1, N2**, and **N3**). Instead, a wide range of *E*_0_ evolved, without noticeably affecting the performance of the evolved networks (Appendix F, Figure A9). This is in line with theoretical considerations (Appendix A), which show that the linear networks (**N1** and **N2**, and the OPR) can evolve optimal performance for *any* value of Interestingly, our individual-based simulations did not converge to arbitrary values of *E*_0_. *E*_0_ ; instead, values in the interval 0 ≤ *E*_0_ ≤ 0.5 evolved in the majority of simulations (Appendix F, Figure A9). We do not entirely understand why the evolution of *E*_0_ should lead to values between 0 and 0.5 and conclude that mathematical expectation alone is not always enough to understand and predict evolutionary outcomes (see also section 4.1.6), even in systems as simple as the ones we used in our simulations.

#### 4.1.6 Network complexity is not a good predictor of performance in the simple learning task

In this section, we considered six types of networks that vary in complexity. They could have one or two layers, a linear or a non-linear activation function and either have an evolving bias or no bias at all. The question arises whether there is a clear-cut relationship between network complexity and network performance.

Figure 6 shows, for each of the six types of networks, the performance of 20 populations of networks that had evolved independently for 500K generations under the conditions of the simple learning task (*S* = 10 and *E*_0_ = 0.5). A darker colour indicates a better performance than a lighter colour. It is obvious that networks with a bias (top) achieve a higher performance than the corresponding networks without a bias (bottom). We do not think that the bias in itself is responsible for this difference. As we have seen in Fig. 5C, bias-free networks can also evolve optimal performance if the initial estimate *E*_0_ is free to evolve. This suggests that having an additional evolutionary degree of freedom (be it the bias or *E*_0_) is what allows networks to evolve an optimal performance.

**Figure 6:**
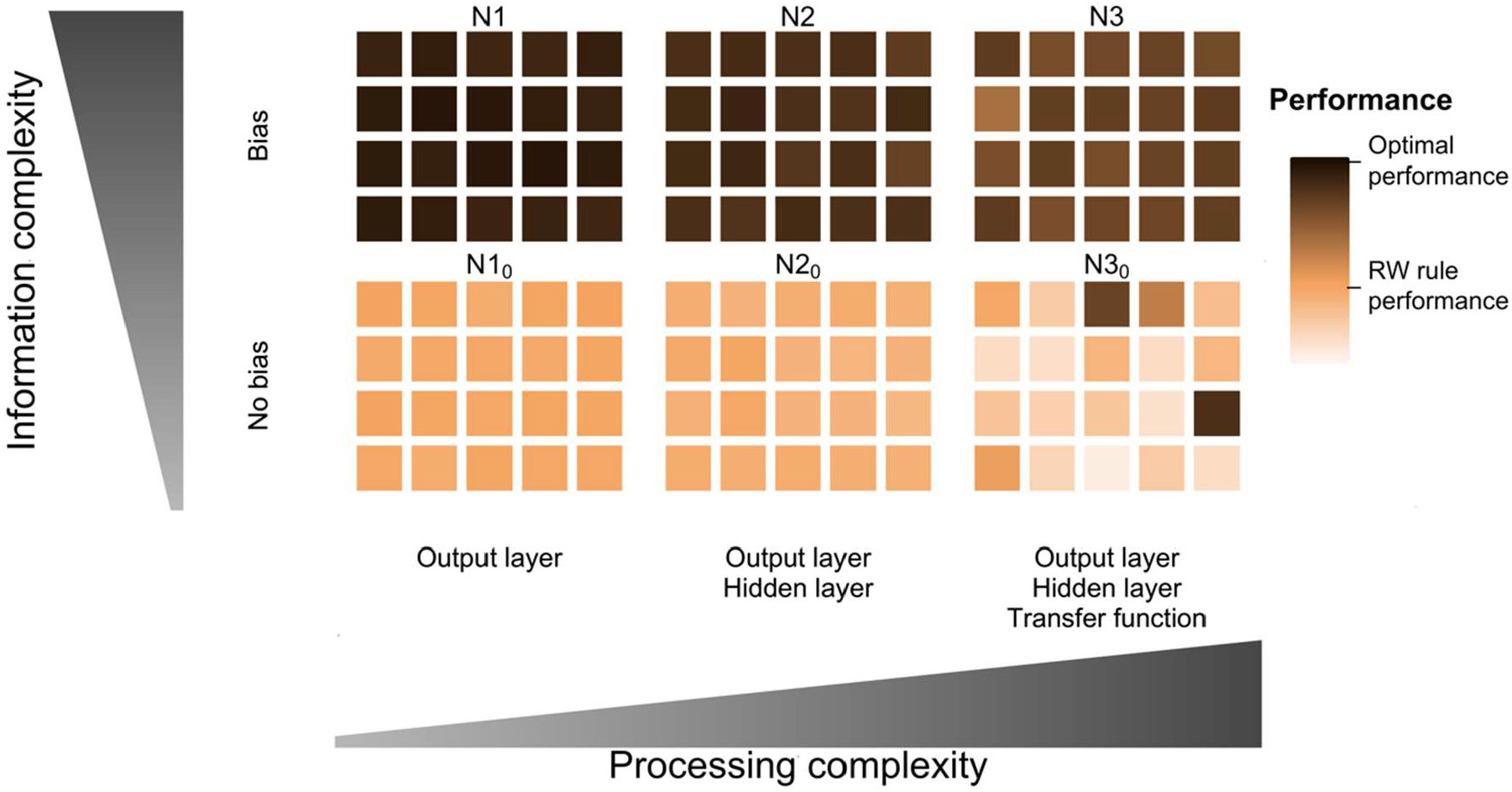
Relationship between network complexity and network performance in the simple learning task with a fixed number of updates and *E*_0_=0.5. Each small square represents a single replicate (20 for each network type) and its colour shows the population mean error over the last 1000 generations of the individual-based simulations that were run for 500K generations.

In contrast, increasing complexity by adding a non-linear activation function usually had a detrimental effect, resulting in worsened performance compared to networks with linear activation function, with the exception of a few replicates in which N3_0_ achieved better performance than N2_0_ (Figure 6 and Appendix C, Figure A2).

The effect of adding a second (hidden) layer to 1-layer networks is more difficult to interpret. On the one hand, it could be said to be negative because, having more weights implied that evolution was slower, and that mutation-selection equilibrium decreased average population performance. On the other hand, this slower evolution could be advantageous once the network performance is close to its optimum. Due to the stochastic nature of the simulations just by chance over a period of time individuals can encounter several environments that have probability of reward away of the average of 0.5 (e.g., skewed towards larger *P* values). Populations with 1-layer networks could rapidly adapt to these environments, which would mean that they would transiently be more severely maladapted when conditions change back to less skewed sequences of environments. For the same situation, 2-layer networks seem to show some kind of robustness due to their slower evolution: theoretically optimal networks (best over all possible *P* value and combination of reward sequences) are present more consistently in the population. Furthermore, **N2**_**0**_ and **N2** were mathematically equivalent to **N1**_**0**_ and **N1**, respectively, but they could achieve this equivalence by different combinations of weights (see Appendix A and Appendix E, Figure A5). Although we did not investigate this further, it could be expected that this phenotypic redundancy, because of the underlying variability in weights, could enhance evolvability in novel environmental conditions [19,20].

In summary, in the simple task, the complexity of the network is not always the best predictor of its performance. In fact, the networks with the best performance, were those with a level of complexity just above the minimum (**N1**).

Nevertheless, this conclusion might not be very surprising if we acknowledge that so far we have considered a very simple learning task. In the next section we will address a learning task that is more difficult and slightly more realistic.

### 4.2 A more complicated learning task

In this set of simulations, we considered a more complex task of estimating *P* based on a variable number of observations *S*. This can be considered a more realistic case, since in natural conditions it is unlikely that organisms face always the same number of observations to learn from. For a given *P* each observation had a fixed chance (of 0.1) to be the last one, with maximum number of experiences set to 20 (similar to Trimmer *et al*. [13]).

For this task we also considered different initial estimates. Here, we are going to focus on the case when *E*_0_ was fixed to 0.5 and mention other results only briefly.

#### 4.2.1 Additional learning rules and networks

When the initial estimate of probability of reward was set to 0.5, Trimmer *et al*. [13] calculated optimal (truly) Bayesian rule as:

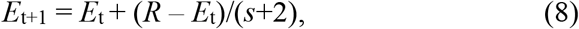

where *s* is the number of experiences that have been witnessed so far, including the current one. This rule is similar to RW rule, but with learning rate that decreases with each observation (1/(s+2)). As Trimmer *et al*. [13] pointed out, the “optimally performing rule” (eq. (5)) yields an optimal estimate only after all updates have taken place but each update is not Bayesian. In contrast, the rule in equation (8) is truly Bayesian, giving optimal estimate at every update. Both rules also show different behaviour (compare Figures 4 and 8 and see Appendix B). We used learning behaviour and error of Bayesian rule and the best performing RW rule as benchmarks when comparing the behaviour and performance of NNs (see section 3.5 and Supplementary Material for more details).

**Figure 7.**
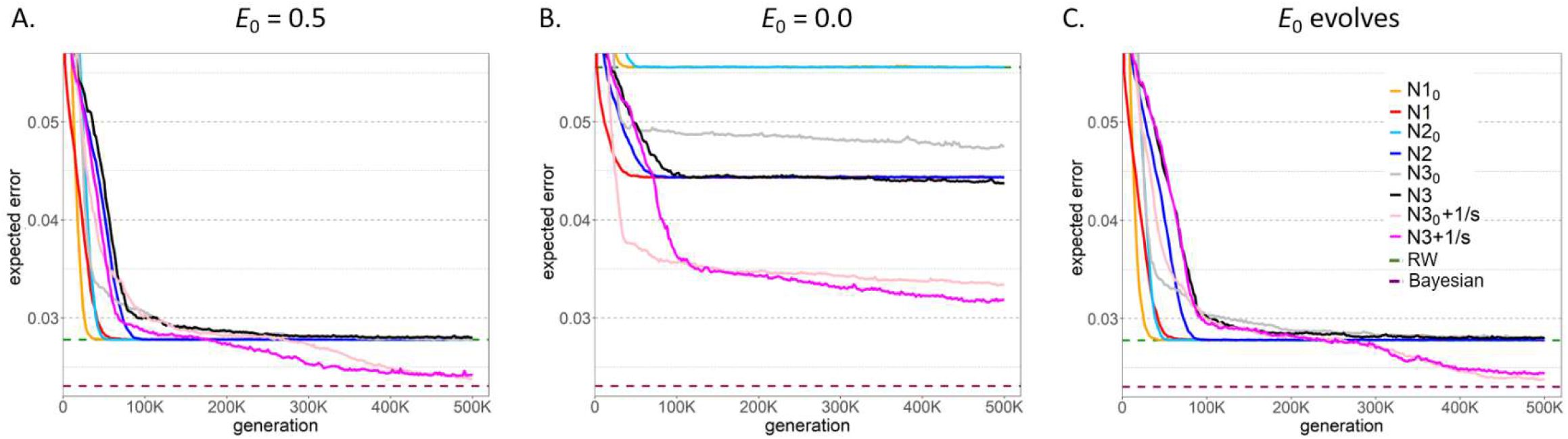
Change in performance of different networks when the number of updates is variable. **(A)** E_0_ = 0.5, the reference networks reach RW Rule performance. Only non-linear networks with additional input (**N3**_**0**_**+1/*s*** and **N3+1/*s***) can reach better performance. (**B**) E_0_ = 0.0, linear networks without bias perform as good as the RW rule. Other networks perform clearly better than it, but optimal performance is not reached within the time limit of the simulation. (**C**) Evolving E_0_ - for all reference networks *E*_0_ evolves towards 0.5 and therefore these networks have performance as in (A), **N3+1/*s*** and **N3**_**0**_**+1/*s*** reach much better performance with E_0_ usually different from 0.5. Linear networks with extra input 1/s are not shown as they do not significantly differ from their two input counterparts. For linear networks average of 10 replicates is plotted, where for each replicate the lowest expected error in the population at the given time was taken into account. Due to error calculation time for non-liner networks only the best replicate is plotted and minimal error is based on a sample of 100 individuals in each generation (sample of 100 gives a good indication of the evolutionary trend – data not shown).

**Figure 8.**
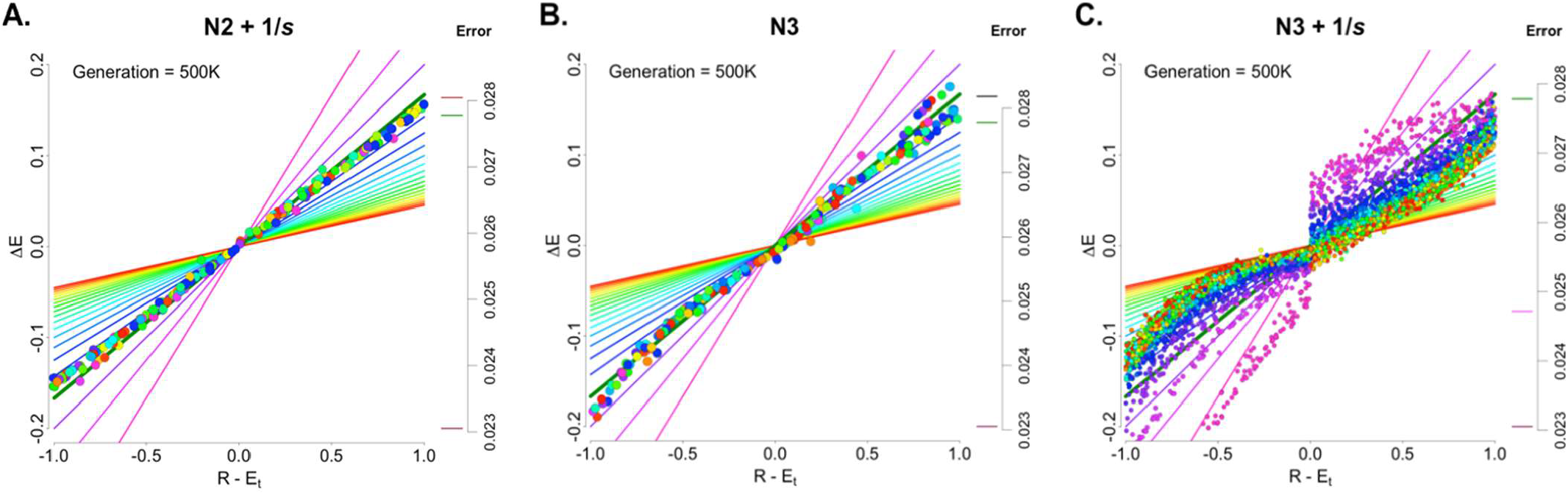
Rescorla Wagner Plots for the networks that evolved in learning scenario with variable number of updates and initial estimate E_0_ = 0.5. Different colours represent data created with different *s* values. Networks **N2+1/*s*** (A) and **N3** (B) show an updating behaviour resembling the RW rule. Network **N3+1/*s*** (C), is able to use the information on update number to display an updating behaviour that, although not identical to that of the Bayesian rule (straight lines in the background), resembles the Bayesian in that early information has larger effect than later information, thus for a given *R-E*_t_ value, change in estimate (Δ*E*) is larger for early updates - small values of *s* (violet end of the rainbow spectrum) and smaller for later updates - big values of s (red end of the rainbow spectrum). This is evident in the clear-cut segregation of data points by colour (that is by *s*), specially for smaller values of *s*, where their effect is larger.

None of the networks considered so far (Figure 2) was provided with direct information of the number of updates. However, if the number of updates varies between environments, the current update number is an important piece of information that could be expected to help the networks to approximate Bayesian rule (eq. (11)) and therefore achieve better performance. For this reason, we also considered networks that resembled previously analysed networks (Figure 2) but had an additional input node, equal to 1/*s*. We will call these networks **N1+1/*s*** … **N3+1/*s***. The extra input initially takes the value of 1 and decreases as the number of updates increases.

Estimating the probability of reward is an intrinsically more difficult task if the number of updates is variable, which is evident in the fact that all rules and networks have worse performance than in the simple learning task. Even the (optimal) Bayesian rule for the more complicated task has a larger error than the RW rule in the simple task (compare scale of Y axes in Figures 5 and 7).

For initial estimate of 0.5, in the task with a variable number of updates all the networks of the previous section converge to behave and perform as good as or close to the RW rule (Figure 7A). Adding an extra input for update count to the linear networks does not improve their performance (see Appendix F, Figure A10). Non-linear networks with additional input (1/*s*), **N3**_**0**_**+1/*s*** and **N3+1/*s***, are the only ones that perform clearly better than RW rule and the rest of the networks, although they never reached optimal performance in our simulations. In the following section (4.2.2) we explore this in more depth.

When *E*_0_ is fixed to 0, all the networks perform clearly worse than for E_0_=0.5 (Figure 7B) and when the E_0_ can evolve for most networks *E*_0_ converges to 0.5 and their performance is identical to E_0_ fixed to 0.5 (Figure 7A and 7C). In this case, when the time available for reaching the final estimate is unknow, initial estimate that is close to the statistically expected *P* value (0.5) seems to be best for most networks and therefore their evolved initial estimate is close to 0.5. Only non-linear networks with 1/s as input evolved different initial estimates and reached performance closer to the optimal one (Figure 7C, Appendix F, Figure A12).

#### 4.2.2 Observing learning behaviour complements understanding

In order to better understand the difference between the best network (**N3+1/*s***) and networks simpler by only a single feature: **N3** (the same as **N3+1/*s*** but without the extra input) and **N2+1/*s*** (the same as **N3+1/*s*** but without the non-linear activation function), we also looked at their behaviour (Figure 8).

In line with what could be expected from their respective performances (see Figure 7A and Appendix F, Figure A10), only **N3+1/*s*** and **N3**_**0**_**+1/*s*** networks showed a behaviour that was clearly different than RW rule and reminiscent of that of the Bayesian rule, i.e., they used information of the number of updates elapsed, to modulate the amount of change in the estimate of probability of reward (Figure 8C).

The fact that the linear networks with additional input (**N2+1/*s***) did not show this kind of behaviour highlights the fact that networks are not always able to efficiently use extra information, even if it seems useful. It is only the combination of both features, the additional information of the additional input 1/*s* and the tanh activation function, that enables networks to surpass RW rule and get closer to Bayesian, not only in performance but also in behaviour. Interestingly, also **N3+1/*s*** and **N3**_**0**_**+1/*s*** networks underwent a stage of their evolution when they were performing and behaving similarly to the RW rule, with little effect of the current update number on their behaviour. Only at later stages they evolved to meaningfully use the additional input information and improve their performance.

It should be noted that whereas the Bayesian rule performs a product of the (diminishing) learning rate and the discrepancy between reward and expected value, the networks are not able to perform product operations directly. Instead, the extra information has to undergo a more indirect and complex transformation during the operation of the network. This in part may explain why these networks approximated, but did not fully replicate the behaviour and performance of the Bayesian rule. More complex networks with multiple processing layers with non-linear activation functions should, in theory, emulate the Bayesian rule and achieve optimal performance [18]. However, the network complexity and evolutionary time needed to accomplish this is not clear.

Investigating the behaviour of different networks allowed us to also notice that performance equal to RW rule does not always guarantee a behaviour that is like it. Contrary to all the cases considered so far, when initial estimate E_0_ was fixed to 0 in the task with variable number of updates, all networks that outperformed RW Rule never showed a behaviour that resembled it (straight line through origin), even during the stage in which their performance was corresponding to that or RW Rule.

#### 4.2.4 Broader comparison of the relationship between complexity and performance

Unlike what we observed for the simple learning task, for the more difficult task, a slight increase in complexity of the networks did not result in improved performance. Intermediate levels of complexity were not enough, only the most complex networks studied by us were able to evolve beyond RW performance (Figure 9).

**Figure 9.**
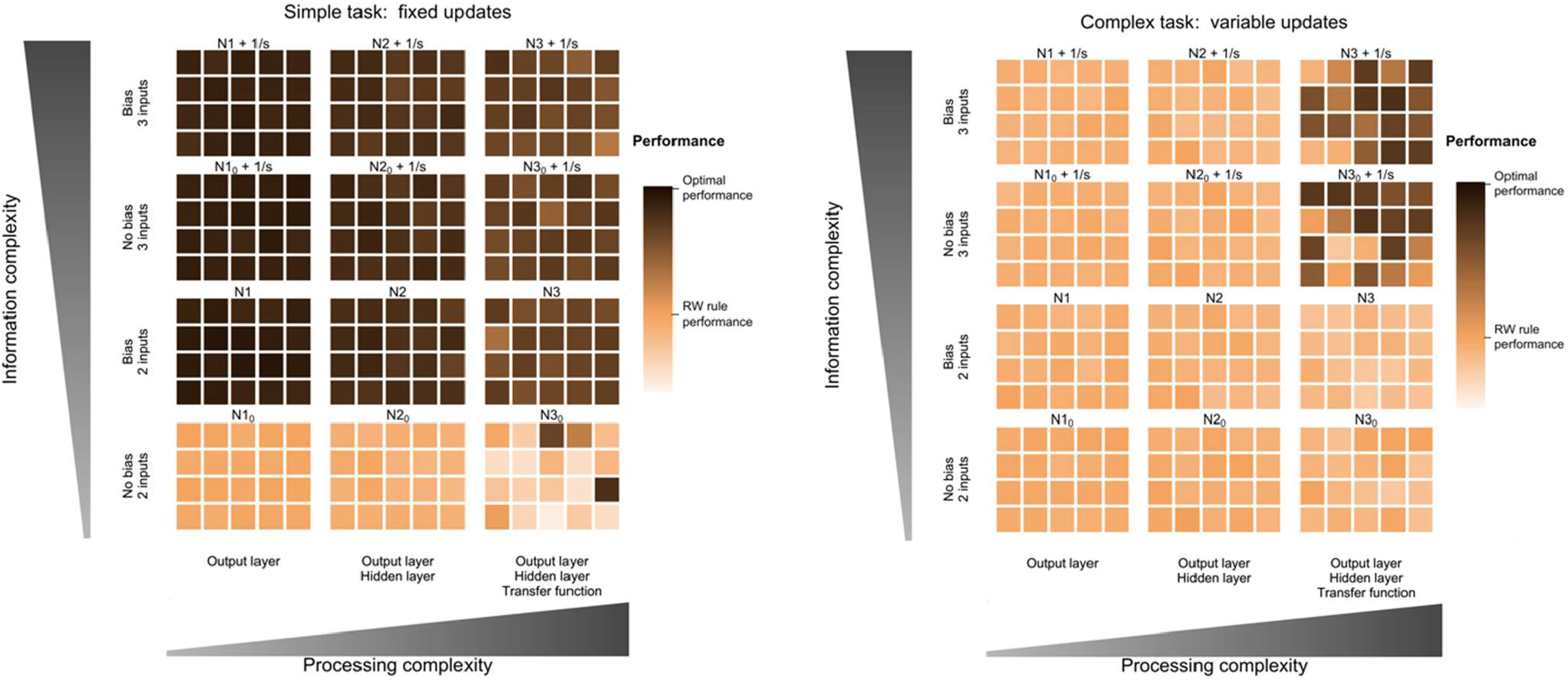
Relationship between network complexity and network performance in both learning tasks and *E*_0_=0.5. Each small square represents a single replicate and its colour shows the population mean error over the last 1000 generations of the individual-based simulations that were run for 500K generations. Performance of the networks is relative to the performance of the rules for a given learning task and it cannot be compared in absolute terms between tasks.

Below we provide a more in detail analysis of the effects of various complexity features on the performance of different networks in both learning tasks.

The very same bias that was beneficial in simple learning task was inconsequential in the more difficult task. And reversibly, hidden layer and non-linearity, which were essential (but not sufficient) for performance improvement in difficult task, were of less impact (N3_0_) or even detrimental in the simple task.

The only feature that was somewhat consistently beneficial was the extra input 1/s. Noteworthy, these beneficial effects of the extra input worked in very different ways for each task. In the complex task it worked together with the activation function to allow the networks to approximate Bayesian performance and behaviour, as explained in section 4.2.3, whereas in the simple task it seemed to act as an extra degree of freedom (akin to the bias or the evolving initial estimate) that allowed the networks to perform and behave similarly to the “optimally performing rule” but with variable, *s*-dependent coefficient (data not shown).

In summary, the relationship between network complexity and its performance is strongly dependent on the learning task. In each task various features of the network can have different effect.

## 5. Discussion

In this study we investigated the evolution of learning using artificial neural networks. While each network did not change during its lifetime, it could be used to improve the estimation of the environmental qualities, based on experiences. If the learning task was simple, many networks (even the simple ones) were able to achieve optimal performance, or at least better than the RW rule that we used as reference. However, some complex networks actually performed worse than the simple ones. Similar conclusions have been reached in a different learning context by [11]. In a more complicated (and more realistic) task, many networks reached RW rule performance, but only more complex networks were able to get closer to the optimal performance. Therefore, complexity needed depends on the nature of the learning task and complexity is not always a good predictor of performance.

Even in such a simple system as the one we developed in this study, we had a few findings that were not in line with mathematical and statistical expectations. For example, all networks with bias and evolving initial estimate are mathematically equivalent because they are all able to perform optimally when the number of updates is fixed, regardless of the initial estimate. Therefore, we would expect initial estimates to evolve to any number between 0 and 1 (or even beyond). However, we saw bias towards lower or higher values (depending on the learning task) in our evolutionary simulations.

We also showed that linear networks with or without processing layer are equivalent from mathematical perspective, yet they differ in evolutionary trajectories. Also, more complex networks allow for larger variability in the evolved mechanism (weights) potentially having higher chance to adapt when the environment changes, increasing their evolvability [21].

Our findings highlight the importance of taking mechanisms into consideration, because they can often have evolutionary effects that cannot be predicted on the basis of pure mathematical analysis. Additionally, different mechanisms may lead to strikingly different results (see e.g. [20,22]). This makes it important to compare different implementations in one study [20,22] or perform studies that check the universality of conclusions of the earlier studies using different approaches (this study). For example, both us and Trimmer have shown that the RW rule seems to evolve readily (at least in the early stages of evolution – see also below), but in our study it was much less often replaced by better rules if the task was more challenging.

Looking at the performance of an evolved network gives only information on how efficient learning is, but not how it is achieved. Studying learning behaviour (how experience actually changes the current knowledge) gives a fuller picture on the evolution of learning. This is especially important when studying complex mechanisms. NNs have multiple parameters and potentially complex structure making it difficult to understand their behaviour by looking at the network weights (and Trimmer *et al*. [13] faced equivalent difficulties when comparing the equations that resulted from their binary trees). We found a straightforward way to compare networks’ behaviour with simple rules and potentially experimental data using RW plots. Using RW plots we show that different networks can use the information on the update number in different ways. We could also show that in most cases, networks show updating behaviour that is similar to the rules they are close to in performance at the moment, but this is not always the case. Therefore, basing conclusions only on performance is not always reliable.

We showed how an assessment of learning behaviour can be achieved through relatively simple and straightforward means. Different learning tasks may need different methods, but devising proper methods for assessing learning behaviour and using them routinely will allow a much better understanding of learning mechanisms and their evolution that cannot be realised on basis of studying performance alone.

Just as Trimmer *et al* [13], we found that most of the networks perform and behave like the RW Rule at least during some stages of evolution. This suggests that RW rule can be so prominent because different underlying mechanisms/architectures promote similar observed/behaviour response. Nevertheless, some other of our findings point to weaknesses of the RW Rule that put into question the assertion that RW Rule is favoured by natural selection. Noteworthy, RW performed very poorly compared to most evolved networks when initial estimate was equal to 0.0, which is the situation actually present in laboratory conditions for which RW was developed (and likely also in nature) since initially neutral stimuli correspond to no initial association.

One possible explanation to this discrepancy is that empirical evidence concordant with RW rule is often observed because natural selection has indeed favoured this “rule”, but not predominantly in the way that Trimmer *et al* [13] have proposed. To begin with, their (and our) study relies on detailed comparison of quantitative data. But we should bear in mind that RW rule was developed to qualitatively (not quantitatively) explain experimental results [12].

More importantly, Trimmer *et al* [13] and us studied the simplification of RW model when only one stimulus is present. However, RW model was developed and gained all its reputation from explaining learning phenomena observed when more than one stimulus is present, such as blocking and overshadowing. Actually, one of the most remarkable features of learning (clearly observed during blocking and overshadowing experiments and that is consistent with the model by Rescorla and Wagner) is that it proceeds in a relatively sophisticated way that would look like causal reasoning, rather than just strengthening of associations based on contiguity of stimulus and reward [23]. It could be argued that having this kind of learning would be highly advantageous.

The open question is: are quantitative models of evolution of simple RW rule useful to explain the success of the full multi-stimuli RW model in experimental psychology? Maybe when talking about evolution of learning we should rather focus on more complex scenarios seen both in nature and laboratory (see below). In particular, to fully understand the selective advantage of RW model, learning situation where more than one stimulus is present have to be considered.

For studies concerning the evolution of behaviour NNs seem to be especially relevant as they allow for a broad range of responses to evolve and also the same responses to be achieved in different ways. This allows for much wider flexibility in possible outcomes of evolution that in most of the analytical studies (but see [10]) while at the same time it provides a relatively straightforward and intuitive framework for linking environmental inputs to behavioural outcomes. However, much more work is needed to fully understand the evolution of learning mechanisms. The ecological scenario considered in Trimmer *et al*. [13] and our study was “deliberately very simplistic to allow for comparison of RW rule with optimal rules [and NNs] of the same complexity and also to avoid confusion with decision-making systems” [13]. By evolving NNs in a more complex environment which includes compound stimuli, and allowing more flexible evolution of the structure of ANNs, we could in the future build a more complete evolutionary picture on the types and quality of rules/mechanisms that allow associative learning. Additionally, animals may not directly estimate probability of reward. There are likely other layers of behaviour, for instance decision making, that could be more important for fitness than only how well the estimated association strength between different stimuli is a reflection of reality.

Instead of evolving networks with predefined structure, allowing for evolution of network architecture will shed a light on whether sub-networks linked to different tasks, like assessment and decision making, will evolve. Inclusion of plastic connection between network nodes, akin to synaptic plasticity seen in natural networks, would also add more realism to evolved learning mechanisms. Such plasticity is the defining feature of networks used for machine learning applications. However, in this field predefined and often biologically implausible learning algorithms are used. Nevertheless, an interesting approach was taken by Chalmers [24] – he used a genetic algorithm to evolve a learning rule for weight updating in a simple NNs. He showed that the delta rule (similar to RW rule) evolves and is one of the most successful rules. This learning rule is also used in the field of reservoir computing [25,26] where most of the network collects and process information and only part of the network directly linked to output is modified by learning. Such approaches provide an interesting avenue in the study of the evolution of learning.

Additionally, learning should be tested in more ecologically realistic scenarios that include for example, simultaneous assessment of various stimuli, exploration-exploitation trade-off, competition, and environmental variability [27–30]. Our future line of research on the evolution of learning does consider foraging decisions in a changing world where learning is achieved by modifying part of the neural network, as inspired by the reservoir computing and biological observations (Kozielska and Weissing, in preparation).

## 6. Supplementary Material

Appendices A-F and simulation code can be found online at https://github.com/marmgroup/AssociativeLearning

